# Distributed implementation of Boolean functions by transcriptional synthetic circuits

**DOI:** 10.1101/2020.04.21.053231

**Authors:** M. Ali Al-Radhawi, Anh Phong Tran, Elizabeth A. Ernst, Tianchi Chen, Christopher A. Voigt, Eduardo D. Sontag

**Author notes:** These two authors contributed equally (M. Ali Al-Radhawi, Anh Phong Tran).

## Abstract

Starting in the early 2000s, a sophisticated technology has been developed for the rational construction of synthetic genetic networks that implement specified logical functionalities. Despite impressive progress, however, the scaling necessary in order to achieve greater computational power has been hampered by many constraints, including repressor toxicity and the lack of large sets of mutually-orthogonal repressors. As a consequence, a typical circuit contains no more than roughly seven repressor-based gates per cell. A possible way around this scalability problem is to distribute the computation among multiple cell types, which communicate among themselves using diffusible small molecules (DSMs) and each of which implements a small sub-circuit. Examples of DSMs are those employed by quorum sensing systems in bacteria. This paper focuses on systematic ways to implement this distributed approach, in the context of the evaluation of arbitrary Boolean functions.

The unique characteristics of genetic circuits and the properties of DSMs require the development of new Boolean synthesis methods, distinct from those classically used in electronic circuit design. In this work, we propose a fast algorithm to synthesize distributed realizations for any Boolean function, under constraints on the number of gates per cell and the number of orthogonal DSMs. The method is based on an exact synthesis algorithm to find the minimal circuit per cell, which in turn allows us to build an extensive database of Boolean functions up to a given number of inputs.

For concreteness, we will specifically focus on circuits of up to 4 inputs, which might represent, for example, two chemical inducers and two light inputs at different frequencies. Our method shows that, with a constraint of no more than seven gates per cell, the use of a single DSM increases the total number of realizable circuits by at least 7.58-fold compared to centralized computation. Moreover, when allowing two DSM’s, one can realize 99.995% of all possible 4-input Boolean functions, still with at most 7 gates per cell. The methodology introduced here can be readily adapted to complement recent genetic circuit design automation software.

## Introduction

### The TF bottleneck in genetic circuit design

Cells can inherently perform intricate operations that include adapting to a new environment, responding to various stimuli, and building complex constructs such as proteins. This is enabled, in part, by the fact that genes can connect to each other in a circuit-like manner via diverse mechanisms that include regulators such as Transcription Factors (TFs). Hence, it has been long recognized that genetic circuits resemble electronic circuits in their ability to process logical operations [1]. The implementation of synthetic circuits for biological computations inside cells is of particular interest in applications ranging from drug delivery to the engineering of micro-organisms, immunotherapy, and biofuel production [2, 3, 4]. Despite its potential, the rational design of genetic circuits was initially very difficult and labor intensive, as it required striking a precise balance of regulator abundances. In addition, the sensitivity of cells to growth conditions was poorly-understood, and it was difficult to characterize cell behavior [5]. Furthermore, the expansion of the set of TFs with negligible cross-talk that perform reliably under a wide range of conditions has been challenging [6, 7, 8].

Recent biotechnological developments have ameliorated some of these difficulties. Firstly, computer-aided techniques have been developed to automate the design process [9]. Secondly, efficient and reliable TFs can now be designed through re-purposing of the CRISPR associated protein, known as Cas9, by deactivating its endonuclease activity [10]. This catalytically-inactive Cas9 known as dCas9 retains the ability to bind to a single guide RNA (sgRNA). The resulting complex acts as a TF by binding to the region on the genome that matches its sgRNA strand. The bound sgRNA-dCAS9 complex turns the targeted gene off by repressing the adjacent promoter or by interfering with the elongation downstream [11]. This new platform, referred to as CRISPR interference (CRISPRi), has a modular nature that enables a single protein dCas9 to target multiple genes at once by changing the associated sgRNA [12]. CRISPRi has been used to build multi-input genetic circuits that consist of NOT and NOR gates [12, 13].

The CRISPRi technology allows to design an extremely large number of orthogonal regulators based on sgRNAs. A limiting factor is the fact that dCas9 is a shared resource amongst the different gates which needs to be continuously expressed at very high rates, and this leads to high toxicity for the host cells [12]. This problem is known collectively as the *dCas9 bottleneck* and has been the subject of a high level of research activity aimed at reducing dCas9 toxicity [14, 15, 16, 17, 18]. Thus, in practice, the maximum reported number of repressor-based gates per cell in a genetic circuit remains stagnant at 7-8 in both E. coli [9, 19] and yeast [13].

### Distributed cellular computation

The difficulty in designing scalable biological computational systems has prompted the development of distributed approaches in synthetic biology [20, 21]. Bacteria can exercise cell-to-cell communication via diffusible small molecules (DSMs) [22] or conjugation [23]. In our context, the problem of TF bottlenecks can be circumvented by distributing the overall computation among several (colonies of) cells, each of which performs a specific part of the computation, thus allowing the same TFs to be reused, and keeping dCas9 concentrations at innocuous levels. This approach has been demonstrated by the construction of all types of two-input Boolean functions in cells, each containing one gate [24]. More recent designs are able to realize 1-input-1-output identity/NOT functions as building blocks in segregated consortia [25] or via recombinase logic [26]. A 3D cell consortium has been designed to function as a full adder [27].

In this work, we are interested in developing a synthesis framework for *diffusible systems* that exchange DSMs. A minimal unit consists of two components: a “sender” consisting of the enzyme that produces a DSM and a “receiver” that responds to this DSM and turns on a promoter. However, the number of sender/receiver pairs that have been used together remains very small. Similar to crosstalk between TFs, there is the additional challenge of building an orthogonal set of DSMs where each receiver only responds to its cognate signal. Using directed evolution approaches [28], the method described in [29] can be applied to create a set of four orthogonal sender/receiver pairs (C4-HSL Rhl, 3OC12-HSL Las, 3OC6HSL Lux and pCou-HSL Rpa). Efforts to expand the set of the orthogonal DSMs to eight are underway.

The aim of this work is to automate the process of Boolean synthesis for diffusible systems, under constraints on the number of available orthogonal DSM molecules and constraints on the number of gates per cell. In state of the art technology, the feasible numbers are around seven gates per cell and a total number of 4-5 DSMs.

### A novel Boolean synthesis problem

Due the nature of the CRISPRi framework and limitations of the current technology, we assume that the Boolean synthesis problem is constrained to use 2-input NOR (NOR2) gates only. (Note that a NOT gate can be implemented as a NOR2 gate with identical inputs.) NOR gates are universal, in the sense that any Boolean function can be represented via NOR gates [30]. In order for cells to communicate with each other, DSMs are used in our design. A cell can release one or multiple DSMs that can act as inputs to other cells or as overall outputs of a circuit.

Given a Boolean Function (BF) and its full circuit representation, the most natural “topdown” approach is to apply graph partitioning algorithms that, in essence, cut the full design into smaller subdivisions [31, 32]. However, these partitioning methods tend to yield designs that require a large number of cuts in order to fulfill the constraints on the number of allowed gates per cell (and thus, requiring the implementation of a large number of DSMs).

Furthermore, since the small molecules are diffusible, if multiple cells release the same DSM, the released concentrations add up, and the DSM will in effect act to implement an “virtual OR gate”. If a cell has a DSM input, then the logical value of that input is the sum of all the logical values of the outputs of all other cells that release that DSM. These particularities call for an alternative approach.

In this work, after introducing a formal mathematical formulation of the problem, we propose an alternative “bottom-up” approach that builds up from the synthesis of individual cells. The next step is to take advantage of the additive properties DSMs in order to compute the output as a disjunction, thus limiting the number of DSMs required. We also propose a graph partitioning algorithm to complement the aforementioned method and create a comprehensive optimized and automated design framework.

Our work uses elements of circuit design theory, combined with the adaptation of a branch-and-bound algorithm originally developed in a different context by one of the authors [33] to build a database of optimal NOR2 representations for all Boolean functions of up to 4 inputs. In comparison, the Cello software [9] produces NOR2 realizations via popular packages such as ABC [34] and Espresso [35] which are not guaranteed to be generate circuits which are optimal (in the sense of having minimal number of gates) [36].

The database as well as the code that is used to generate the various designs in this work are made available through GitHub.

### Problem Statement

Let *n* be the number of inputs, and *p* be the number of outputs, such as reporter signals. We are given two positive integers *N, q*, which are the maximum number of NOR2 gates per cell, and the maximum number of orthogonal DSMs, respectively. Mathematically, a vector valued Boolean function (BF) *F*: {0, 1}^*n*^ → {0, 1}^*p*^ can be thought as a vector (*f*_1_,.., *f*_*p*_) of scaler-valued BFs *f*_*j*_: {0, 1}^*n*^ → {0, 1}, *j* = 1,.., *p*. Each BF can be succinctly represented via a hexadecimal code (see Methods). Given *N* and *q*, we are interested in the problem of finding a distributed circuit realization that obeys the two following constraints:

1. each cell is constrained to use a maximum of *N* NOR2 gates,
2. a total of no more than *q* DSMs are allowed,

If a realization exists that satisfies conditions 1 and 2, then *F* = (*f*_1_,.., *f*_*p*_) is said to be (*N, q*)-*realizable*. A mathematically precise definition is provided in the Methods section.

## Results

Current synthetic biology applications, from the detection of environmental toxins to the design of engineered immune cells, involve typically only a handful of inputs. Thus, even though the methods that we introduce can be, in principle, scaled to an arbitrary number *n* of inputs, for computational tractability we will we restrict our attention to BFs of 4 inputs or less. Note that there are 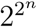 BFs with *n* inputs. This is because there are 2^4^ possible binary strings of 4 bits (0000, 0001, …, 1111), and each of these can be mapped into either a 0 or a 1. In particular, there are 2^16^ possible Boolean functions on 4 inputs.

In the following sections, we discuss the number of functions that can be realized with no DSMs (so that a single colony is sufficient), and then proceed to the topic of this paper proper, namely larger numbers of DSMs.

### (*N*, 0)-realizable networks via exact synthesis

As a first step, BFs that are realizable without any DSMs are considered. We use a branch-and-bound algorithm (see Methods) to find the minimal number of gates required to realize a given BF. The results are summarized in Fig. 1. In particular, we find that only about 11.69% of the 2^16^ different BFs with 4 inputs can be realized using seven NOR gates or less. Moreover, we find that 4-input BFs can require up to fourteen NOR gates, and they most frequently require ten NOR gates, with a median of 10. Thus, most realizations are not implementable in a single cell via the current technological limitation of around seven gates per cell.

**Figure 1:**
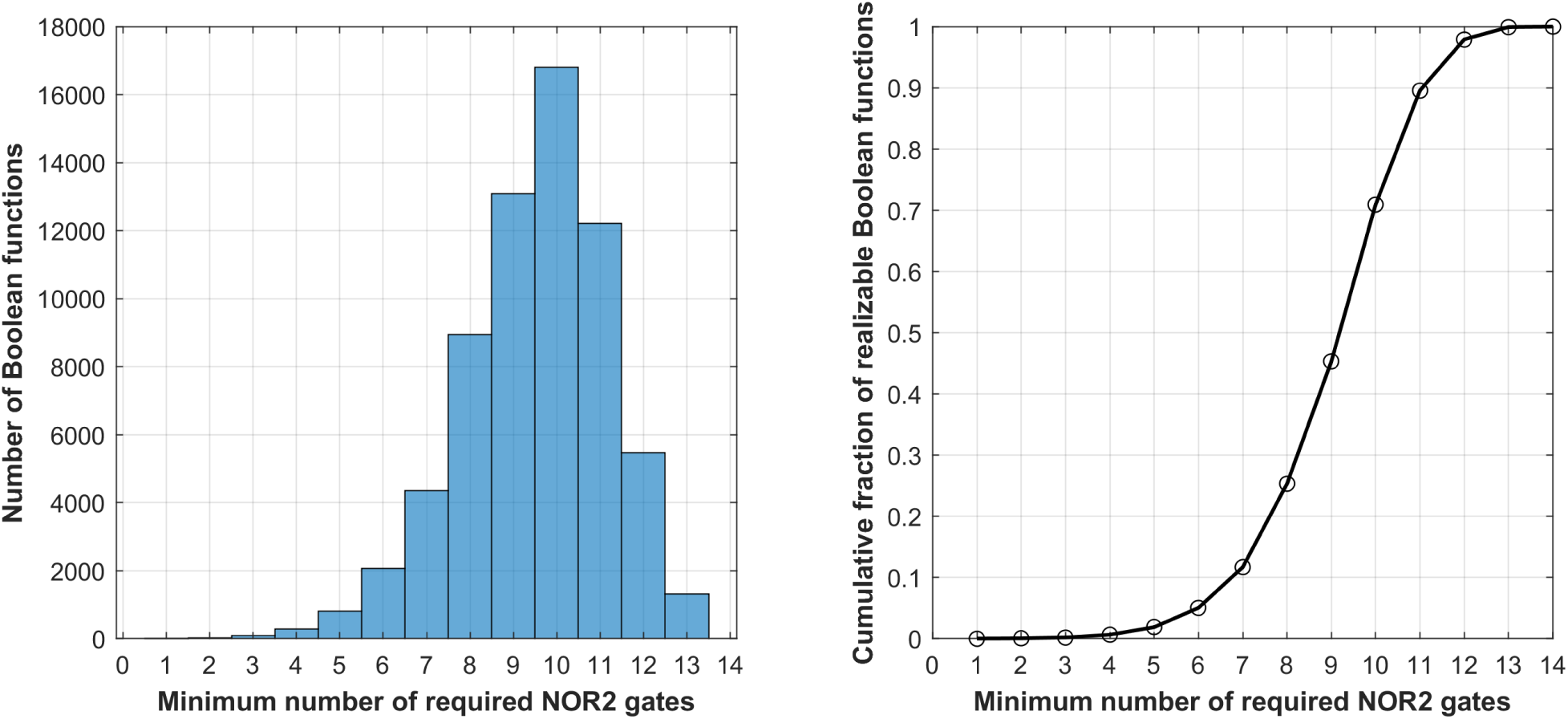
The minimal number of gates required to represent a given 4-input Boolean function. This assumes no use of DSMs, or in the language of synthetic biology, circuits that can be built within one cell. (a) Total number of BFs vs. minimal number of gates needed for realization, (b) the cumulative fraction of realizable Boolean functions vs. the number of NOR2 gates.

The algorithm was used to generate the data for the histograms in Fig. 1, but the main goal was to populate a database that provides the optimal (minimal number of gates) realization for every possible 4-input function. This database is used subsequently as a lookup table, to assign an optimal NOR2 design per cell when building distributed designs.

### (*N*, 1)-realizable BFs via disjunctive form design

If a circuit design cannot readily fit in a cell (or equivalently, it is not (*N*, 0)-realizable), we can use a single DSM to distribute the computation. Several designs are possible. We propose a design depicted in Fig. 2 to realize a disjunctive form of *f*. Each cell releases the same DSM. The total concentration of the DSM adds up, hence its concentration acts as an OR gate. Hence, BFs *c*_1_,.., *c*_*m*_ are to be found such that

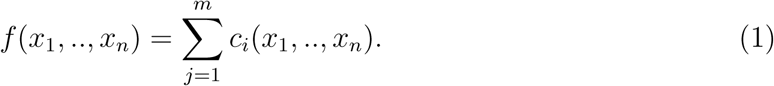

**Figure 2:**
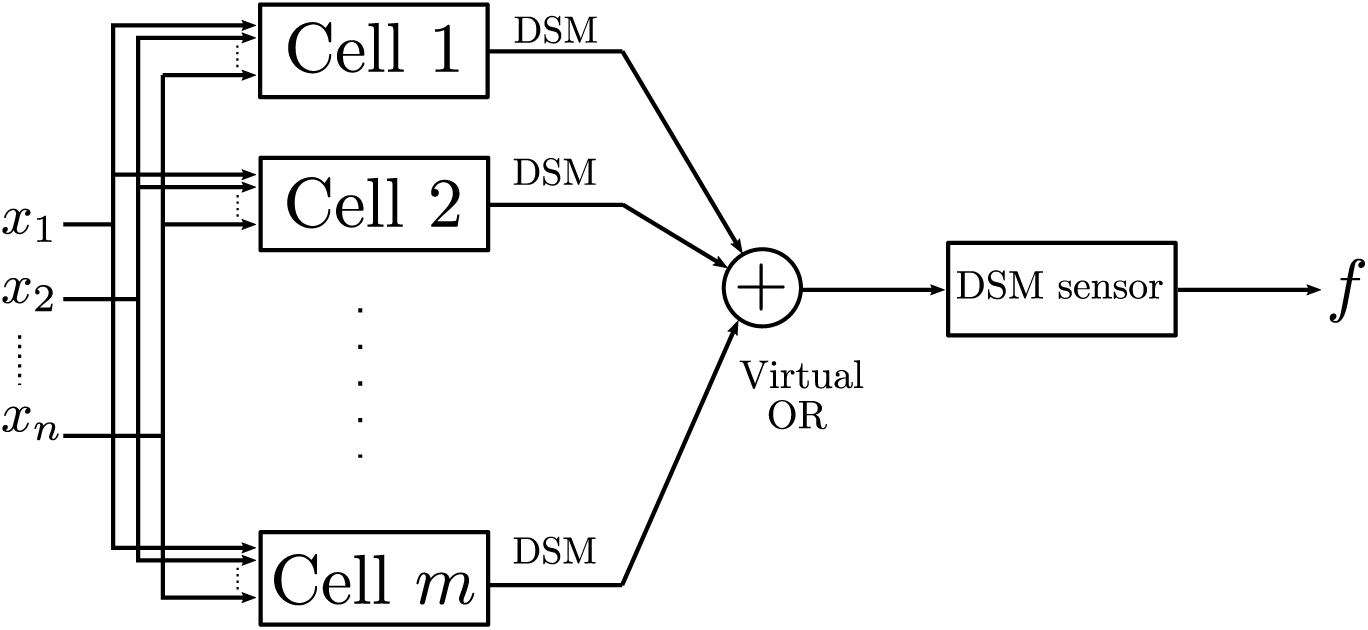
An (*N*, 1)-realization. The figure depicts a distributed computation design for implementing a Boolean function *f* of *n* Boolean variables with one DSM. A DSM sensor can be a DSM-to-GFP cell for instance.

(We interpret the summation in the Boolean sense, as as OR gate.) The summation in (1) uses one DSM. If all the BFs *c*_1_,.., *c*_*m*_ can be realized with *N* gates or less then *f* is (*N*, 1)-realizable. Such a design implements some of the computations outside the cells, and this allows us to realize functions that cannot be realized via graph partitioning with a single cut (see Fig. 4 as an example).

The next step is to determine the BFs *c*_1_,.., *c*_*m*_. One method is to write *f* as a minimized disjunctive form (MDF) which implies that *c*_1_,.., *c*_*m*_ are *product terms* (see Methods for definition). Hence, each cell will compute the corresponding product term in the MDF of *f*. This method allows us to compute the number (*N*, 1)-realizable functions that can be written in the form (1). Furthermore, this procedure produces a *modular* design since *minterm-computing cells* (MCCs) can be designed once and reused as needed in the design of a single or multiple BFs. For instance, a four-input function needs only 13 types of such cells to compute minterms of up to 4 variables. For instance, 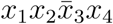 and 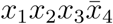 can be computed by the same circuit after permuting the inputs.

For example, when applied to the BF 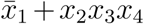, the DNF method will provide a design consisting of two types of cells, one implemented by just one gate and the other one by six gates. While giving a (6, 1) realization, the imbalance between the computational resources allocated to each of the two types of cells may be undesirable in some applications. Thus, we will discuss an approach in which a user is able to choose among different trade-offs between *N* and *q*, along a “Pareto front” of optimal solutions.

#### Realizability test

We provide now a fast test that determines a lower bound on the number of (*N*, 1)-realizable BFs. It relies on development of the concept of *offending minterms*, which are minterms which cannot be realized with *N* gates or less. We use 𝒪_*N*_ to denote the set of all such minterms. For instance,

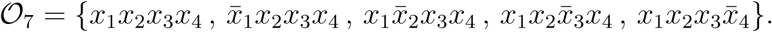

After computing the set 𝒪_*N*_, we show that a BF *f* is (*N*, 1)-realizable if the set of prime implicants of *f* does not contain an offending minterm, and hence a combinatorial test can be developed and is provided in the Methods.

In the case of *N* = 7, the test shows that there are at least 56, 608 (7,1)-realizable 4-input BFs out of a total of 2^16^ = 65, 536 BFs. Therefore, the percentage of realizable BFs is at least 86.384% when using one DSM, compared to only 11.690 % without any DSMs for 4-input BFs, which amounts to a 7.39-fold increase. In the case of *N* = 8, the percentage of (8,1)-realizable functions is at least 96.877%, compared to 25.321% for (8,0)-realizable functions. Note that these bounds are not tight, as we will show in the next subsection.

### Realization with offending minterms

We have shown that if there are no offending minterms in the minimized disjunctive form, then the design shown in Fig. 2 is an effective way to distribute the computation. In this subsection, we generalize the method in order to handle the case of offending minterms. First, we propose a generic method that aims at providing an upper bound on the number of DSMs needed to realize *p* BFs simultaneously. We summarize here the results for the cases of *N* = 7, 8, 9 (which are the given bounds on the number of gates per cell).

#### Theorem 1.

*Given a 4-input multi-output BF F* = (*f*_1_,.., *f*_*p*_). *Then*:

1. *For N* = 7, *there exists* 0 ≤ *q ≤ p* + 2 *such that F is* (7, *q*)*-realizable*.
2. *For N* = 8*, there exists* 0 ≤ *q ≤ p* + 1 *such that F is* (8*, q*)*-realizable*.
3. *For N* = 9*, there exists* 0 ≤ *q ≤ p such that F is* (9*, q*)*-realizable*.

A constructive proof is provided in the Methods section by relying on realizing the offending minterms individually. If an offending minterm appears in multiple BFs, then it only needs to be computed once. For instance, consider the case when *x*_1_*x*_2_*x*_3_*x*_4_ is the only offending minterm that appears in the MDFs of *F* = (*f*_1_,.., *f*_*p*_); then, its 9-gate realization can be partitioned via one DSM to produce two cells whose number of gates just are 4 and 5 (Figure 3-a). More offending minterms can be handled also. Consider, for example, the case when the only offending minterms are 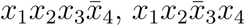. Then, the common product term *x*_1_*x*_2_ can be computed first, and then is communicated to two other cells that compute the required minterms via a single DSM, resulting in a design with a total of 3 cells (with 4 gates each) and one DSM (Figure 3-b). Other cases are described in the Methods.

**Figure 3:**
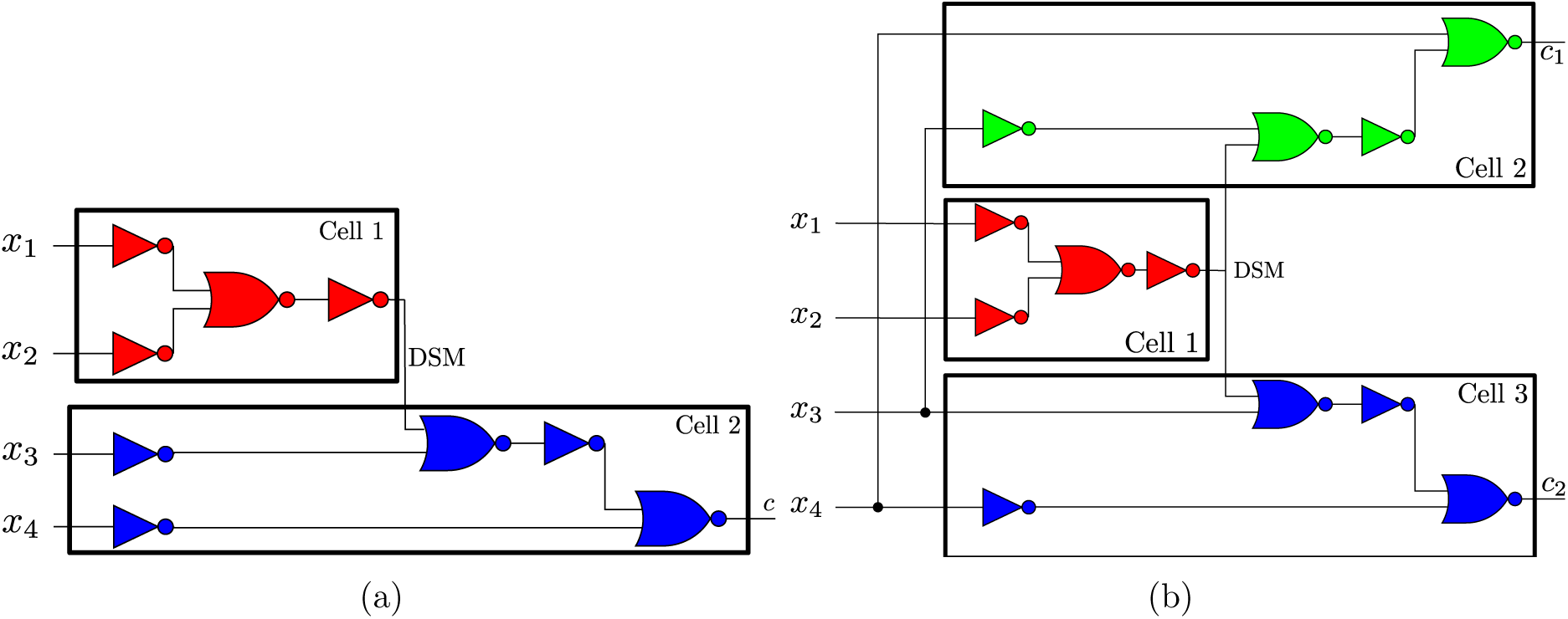
Partitioning of offending minterms. Each diagram can be included as a module in a bigger network that contains the respective offending minterm. (a) Partitioning a realization of the minterm *c* = *x*_1_*x*_2_*x*_3_*x*_4_ that requires a minimum of 9 NOR2 gates. Many partitions are possible. We show here one partition into two cells of 4 and 5 gates respectively, via the use of 1 DSM. (b) Partitioning a realization of two minterms sharing two literals 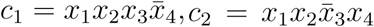. These two minterms that require a minimum of 8 NOR2 gates each can be split up in such a way that the computation for the common product term *x*_1_*x*_2_ can be reused. This allows to use a total of three cells instead of four. Different colors indicate different cells.

#### Scalar BF with offending minterms

The results in the former subsection can be slightly improved in the case of a scalar BF and full databases of realizations with arbitrary *q* can be provided. Furthermore, We provide lower bounds on the percentage of realizable 4-input BFs via the disjunctive form design as shown in Table 1 (see Methods for details). Estimates are also provided for 5-input Boolean functions in Table 2.

**Table 1:** Percentage of realizable 4-input Boolean functions via the disjunctive form design (lower bounds).

| $(N, q)$ | | $q$ | | | |
| --- | --- | --- | --- | --- | --- |
|  |  | 0 | 1 | 2 | 3 |
| $N$ | 7 | 11.690% | 86.384% | 99.951% | 100% |
|  | 8 | 25.321% | 96.877% | 100% | 100% |
|  | 9 | 45.297% | 100% | 100% | 100% |
|  | 10 | 70.939% | 100% | 100% | 100% |
|  | 11 | 89.570% | 100% | 100% | 100% |

**Table 2:**
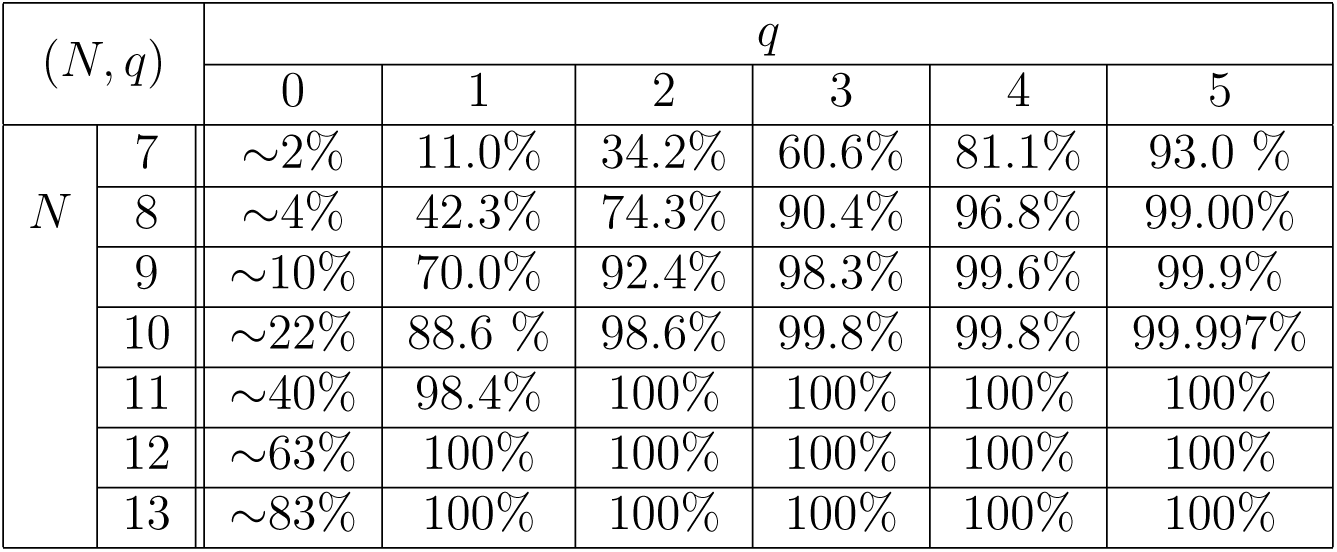
Estimates of the realizable 5-input Boolean functions via the disjunctive form design (lower bounds). The (*N*, 0) percentage values were estimated from the limited list of calculated optimal designs of 5-input Boolean functions provided in [33]. The (*N, q* > 0) values were estimated by examining 100,000 DNF decompositions picked at random from the set of 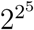 possible 5-input combinations. The reductions due to redundancy when dealing with more than 1 offending minterm were not accounted for, unlike the 4-input case. Thus, the listed estimations of the lower bounds are expected to be looser.

### A graph partitioning algorithm

There are some cases in which the number of DSMs needed can be reduced with respect to the disjunctive design. We propose a graph partitioning algorithm applied to a full NOR2 network generated by the exact synthesis algorithm. In this formulation, the usage of a single DSM will be equivalent to a single cut in the extended NOR2 graph. See Methods for a detailed description of the algorithm.

Due to the nature of graph partitioning, unless the circuit readily fits inside a cell, the use of a DSM signal is necessary. Thus, for 4 inputs and *N* = 7, 11.690% of BFs can be realized without a DSM, another 37.680% of BFs can be realized using 1 DSM, and an additional 34.900% using 2 DSMs, using this technique. This leaves 15.730% of BFs that cannot be realized using 2 or less DSMs. Thus, given the fact that the number of mutually orthogonal DSMs is considered to be a limited resource, the disjunctive-form design is generally preferable as a way to minimize the use of DSMs. However, there exist cases for which the partitioning algorithm offers a more compact realization.

By combining both the disjunctive form design and graph partitioning, the numbers presented in Table 1 can be improved (for the case *q* = 1) to those shown in Table 3.

**Table 3:** Lower bound on cumulative percentage of realizable 4-input Boolean functions combining the disjunctive-form design and graph partitioning algorithms

| $(N, q)$ | | $q$ | | | |
| --- | --- | --- | --- | --- | --- |
|  |  | 0 | 1 | 2 | 3 |
| $N$ | 7 | 11.690% | 88.666% | 99.951% | 100% |
|  | 8 | 25.321% | 97.440% | 100% | 100% |
|  | 9 | 45.297% | 100% | 100% | 100% |
|  | 10 | 70.939% | 100% | 100% | 100% |
|  | 11 | 89.570% | 100% | 100% | 100% |

### Optimized distributed design framework

The designs presented so far provide a circuit representation that fulfills the physical constraints imposed (on numbers of gates and cell types), but requires the use of at least as many cell types as there are minterms in the DNF form. Using the database developed through the exact synthesis algorithm, it is straightforward to test all the combinations of various non-offending product terms that are (*N*, 0)-realizable. This leads, in most cases, to a reduction in the number of required cells for a given realization.

In general, there is a trade-off between the maximal number of gates per cell (*N*) and the number of DSMs (*q*). Depending on the application, different optimization criteria may be suitable. The most intuitive criterion is to use as few cells and NOR gates as possible. In other cases, one may want to minimize the number of cells required, while also minimizing the standard deviation of the number of NOR gates required per cell, in order to reduce differences in computational time between cells.

The use of a look-up database allows for a rapid and exhaustive search of these optimal designs. This is possible by taking all the non-offending minterms and testing for all possible combinations of a given set of minterms. For example, the five possible combinations of the set {1, 2, 3} are {{1, 2}, 3}, {1, 2, 3}, {{1, 3}, 2}, {1, {2, 3}}, and {{1}, {2}, {3}}. In this case, there are five different ways of combining 3 minterms. Given the use of the database, the computation required to compute all five designs is very minimal. This allows a rapid look-up of what minterms can be efficiently combined, while also meeting the constraints of the problem.

Finally, the partitioning algorithm can also be combined with the distributed disjunctive-form algorithm through combining offending and non-offending minterms to find combinations that lead to more compact or desirable designs.

### Illustrative examples and Pareto trade-offs

Various circuit realizations generated by the developed algorithm are provided for different BFs under the constraint that *N* = 7 are shown in Figs. 4, 5, and 6. In Fig. 4(a) and (b), two (7, 1)-realizable circuits are shown with both circuits having six and seven product terms in their DNF form. However, given the look-up table, these realizations are reduced from requiring six and seven cells in the disjunctive-form design to simply requiring two and three cells, respectively. Through combining various minterms, it was found that it is not possible to decrease the maximum NOR gates requirement. In other words, if a minterm is offending, it was not possible for us to find a combination of this minterm with the other minterms of a given BF to realize it with a smaller number of NOR2 gates. However, through these examples, we illustrated that it can be quite straightforward to reduce greatly the required number of cells through the exhaustive search of the optimized disjunctive-form design.

**Figure 4:**
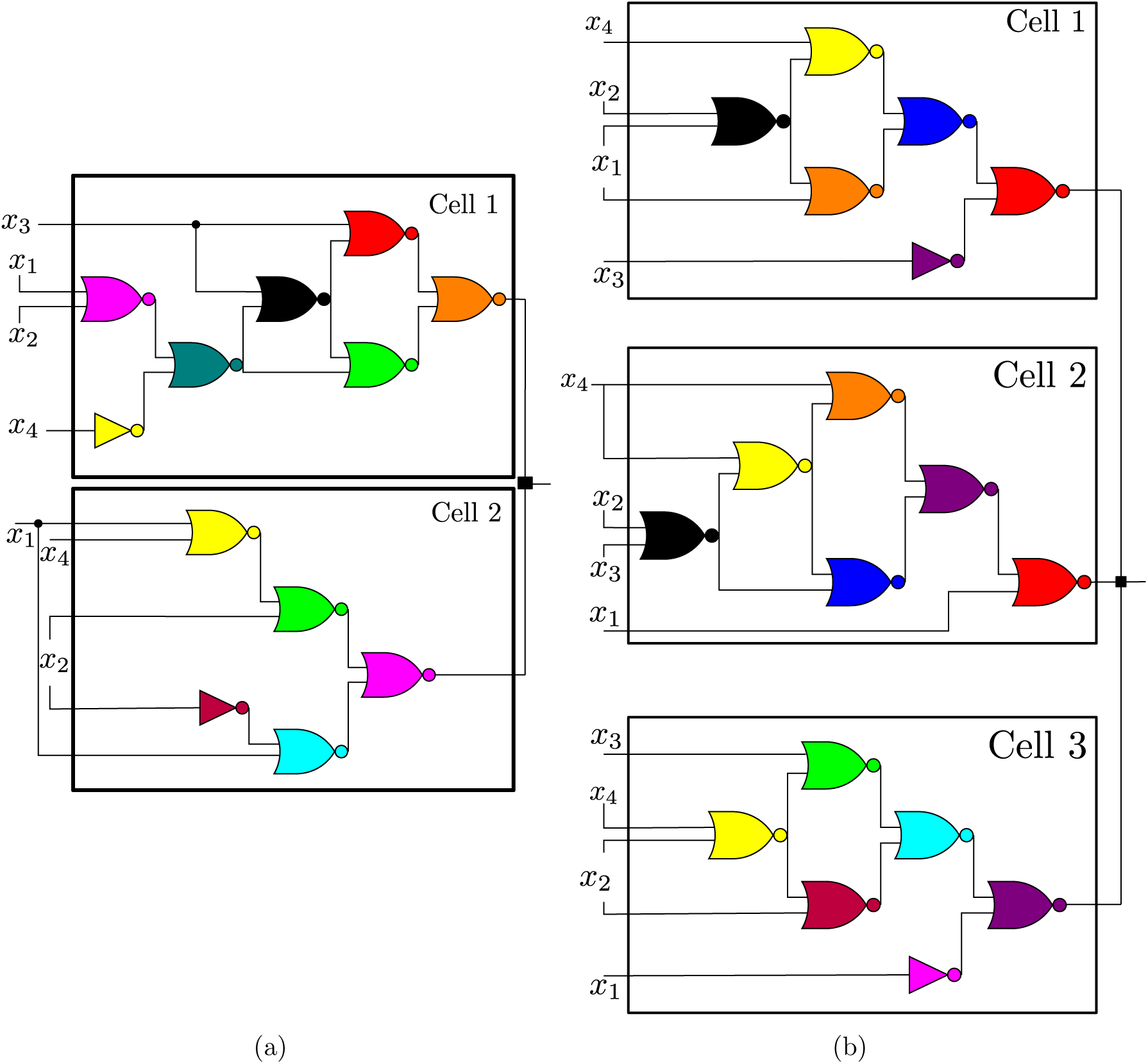
(7,1)-realizable circuits using the proposed algorithm for two Boolean functions. (a) Distributed realization of the Boolean function with the hexadecimal representation (0xE99F), or equivalently 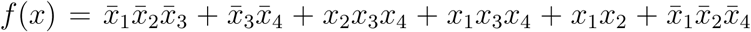, that requires 13 gates to be implemented in a cell. The corresponding graph cannot be partitioned with one DSM. Nevertheless, the optimized disjunctive form provides a design of two cells with 7 and 5 gates as depicted above. This is notable because the distributed design requires less gates than the full circuit using NOR2 gates. (b) The Boolean function (0×977E) requires at least 14 gates to be implemented in a cell. The optimized design yields a design using three cells of 6 gates each. The DNF form is given by 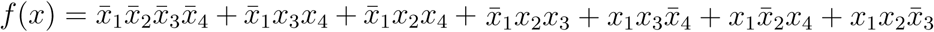. The symbol “■” refers to a DSM sensor.

**Figure 5:**
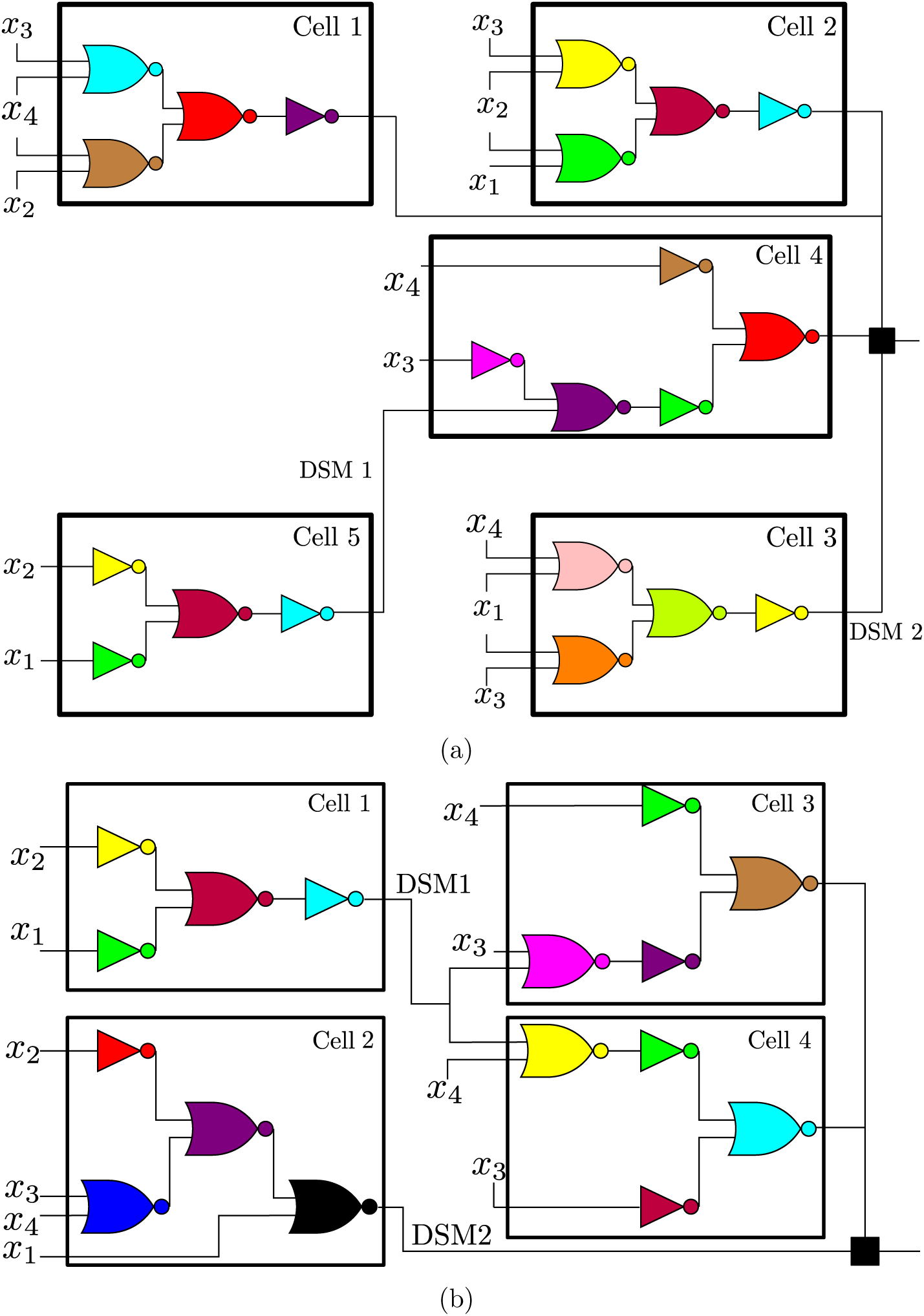
Circuit realizations using the proposed algorithm for two Boolean functions with one and two offending minterms. (a) The function (0xFEE9) requires 14 gates to be implemented in a cell. The DNF form is provided by 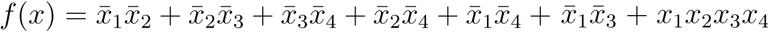. The optimization procedure was used to reduce the number of required cells from 7 (based on the DNF form) to 5, while also ensuring that the final design is (5,2)-realizable. The symbol “■” refers to a DSM sensor. (b) The BF (0xF806) requires 11 gates to be implemented in a cell, and the resulting graph cannot be partitioned with one cut. The DNF form is provided by 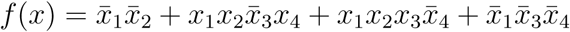. The optimized design required four cells with a unique “sender” (Cell 1) that acts upon two “receivers” (Cells 3,4), making this design (4,2)-realizable.

**Figure 6:**
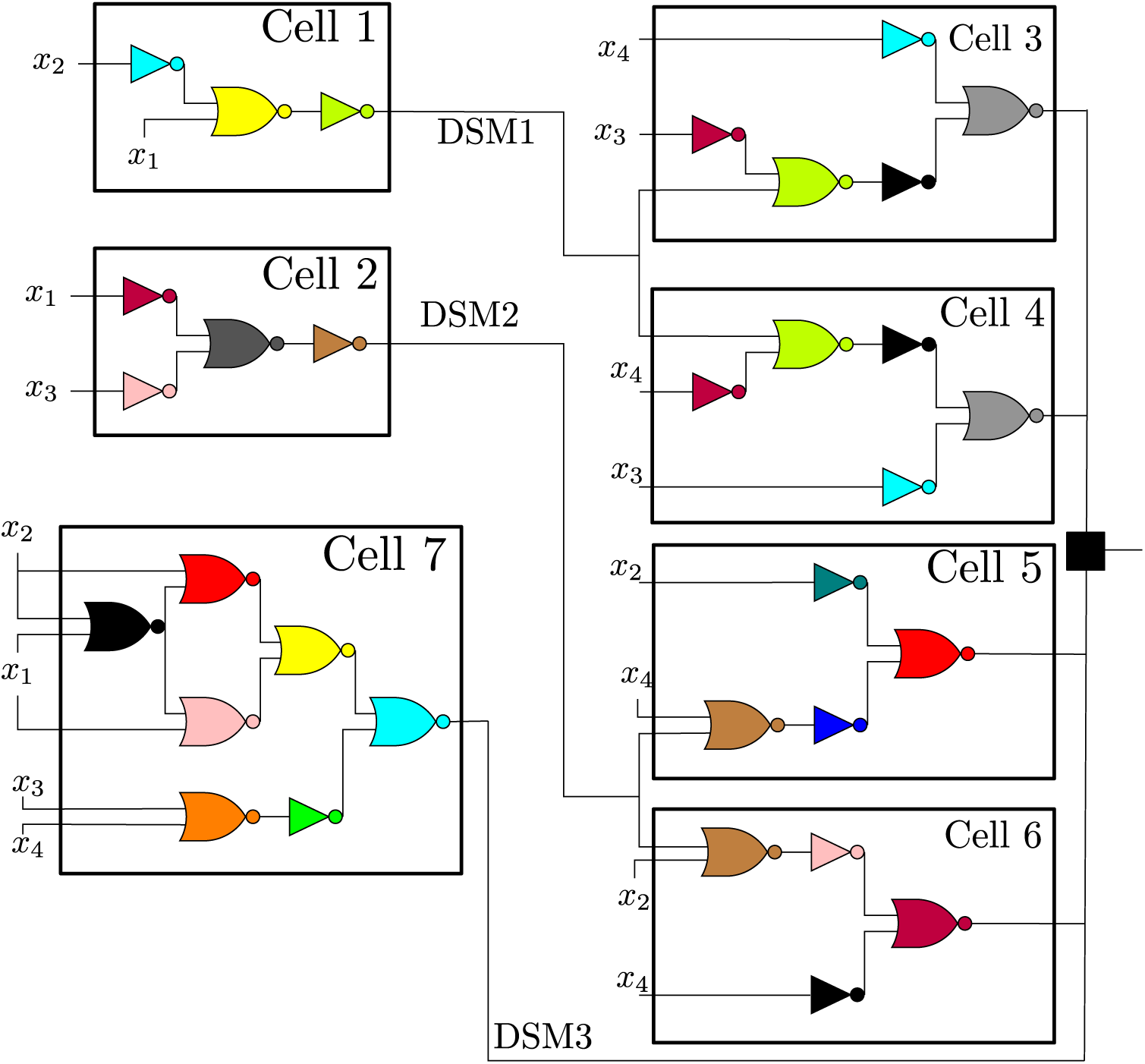
Circuit realizations using the proposed algorithm for a (7-3)-realizable Boolean function. This function (0×2196) requires at least 13 gates to be implemented in a cell. The DNF form is provided by 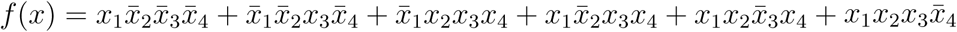. The symbol “■” refers to a DSM sensor.

It is then shown how the algorithm handles offending minterms. Fig. 5 depicts disjunctive form realizations of a BF for which a DSM is necessary to deal with the offending minterm. In Fig. 5(a), Cell 5 is used to create the first DSM signal, which feeds as an input to Cell 4. Cells 1 through 4 combine their outputs through a DSM sensor that acts as an OR gate. Fig. 5(b) shows an example dealing with two offending minterms for which the resulting optimized DNF design is (4,2)-realizable.

In Fig. 6, two different DSMs are used to deal with the three offending minterms. In that design, cells #2 and #3 share a similar input circuit releasing a common DSM. The remaining four non-offending product terms are combined to form two cells requiring seven NOR gates each. This (7,3)-realizable circuit is distributing the computation in two steps by first computing the DSM1 and DSM2 signals from cells 1 and 2, and then computing the operations in cells 3 through 7.

While only a few examples are shown here, the developed algorithm can generate optimized realizations for all 65536 possible 4-input BFs.

Show in Fig. 7 are various designs that can be generated from the DNF form depending on the criteria of selection that can be, for example, the average number of gates per cell, the total number of gates in the design or the range of the gates in final design.

**Figure 7:**
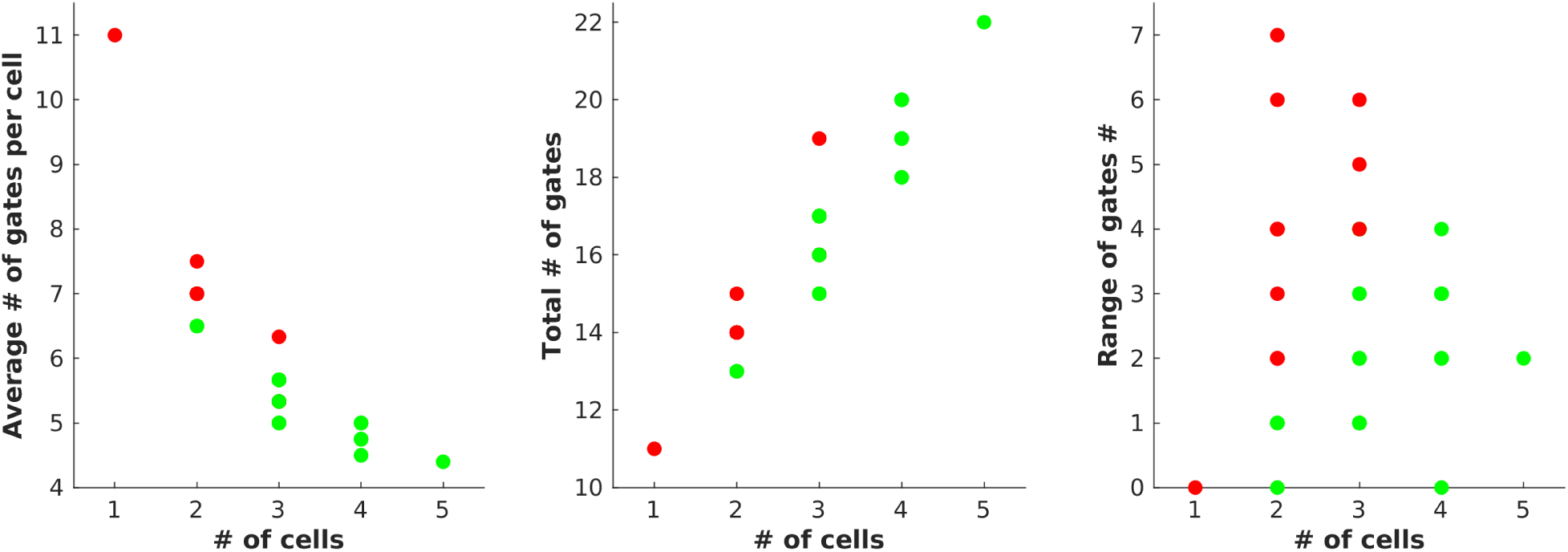
Various possible circuit designs using the exact synthesis library are shown for the non-offending minterms of (0xA7D4). While the focus here was on minimizing the total number of gates, other optimization criteria can be readily implemented. Green bullet points indicate designs that are realizable within the constraint of 7 NOR2 gates per cell, while red bullet points indicates designs which are unrealizable.

In order to illustrate the complementary nature of a unified algorithm that integrates the partitioning and DNF approaches, we show in Fig. 8 how the DNF decomposition of a circuit can yield a (7,1)-realizable design that would not be otherwise possible with a partitioning algorithm. In Fig. 9, the opposite scenario is shown, in which the partitioning algorithm of the full circuit provided by the exact synthesis algorithm yields a very compact (7,1)-realizable design of 2 cells using 5 and 7 gates. A comparable design in the DNF form would require at least 3 cells and 2 DSMs.

**Figure 8:**
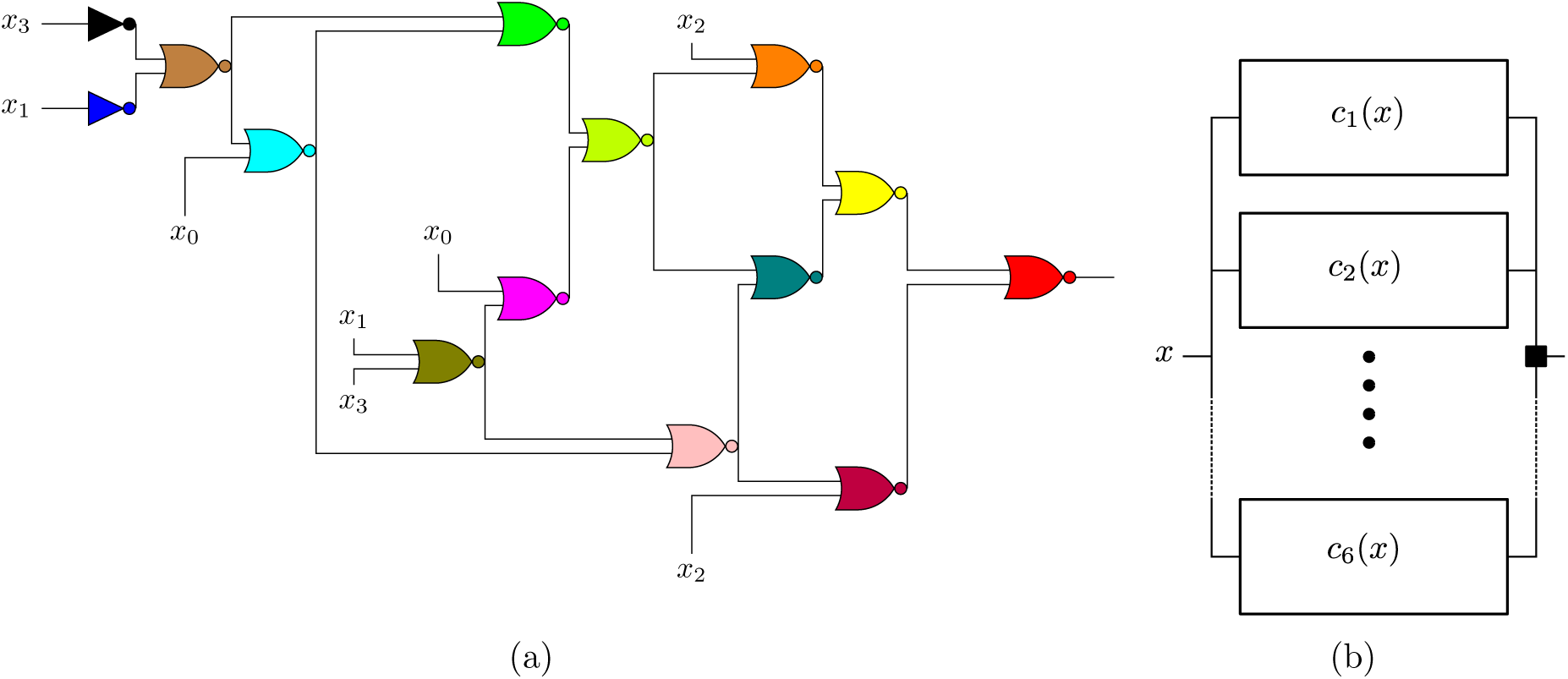
Depicted here is an illustrative example for which the DNF design is a lot more economical in terms of required number of DSMs. (a) The exact synthesis design representing (0×1668) is shown where it requires at least 14 NOR2 gates. There exists no partitioning using 2 DSMs that can make this representation (7,2)-realizable. (b) The proposed disjunctive design re-synethesize the logic from the bottom up and it provides a (7,1)-realization with six cells each containing 7 gates. The BFs are 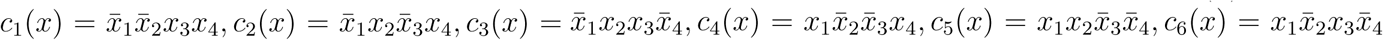. The symbol “■” refers to a DSM sensor.

**Figure 9:**
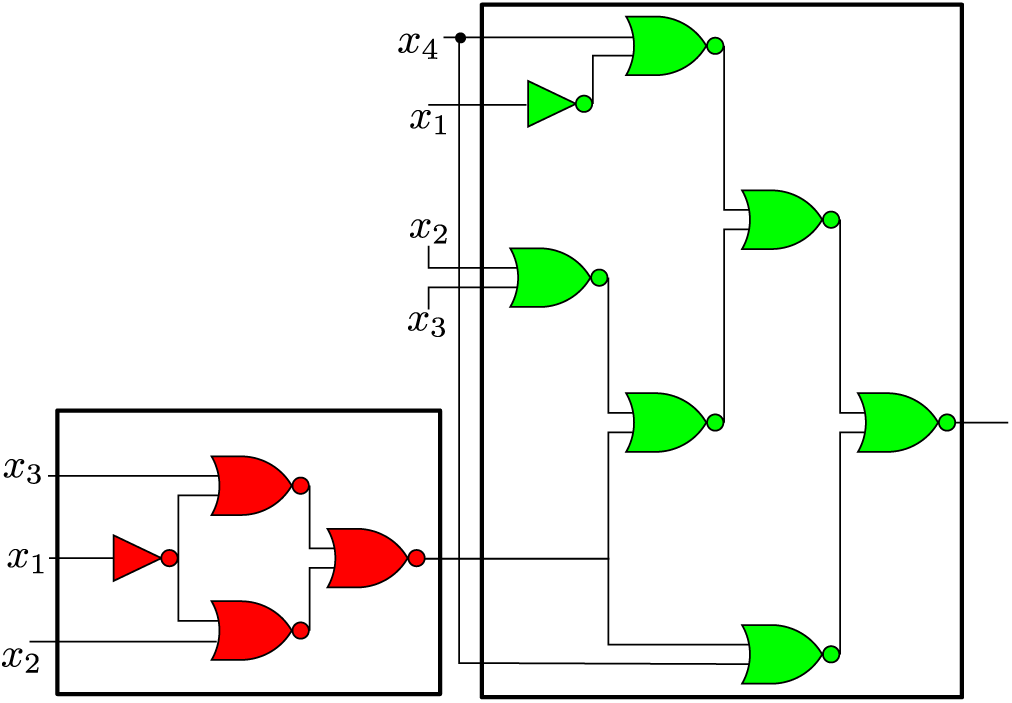
Depicted here is an illustrative example for which the design using the partitioning algorithm is more compact than the most compact DNF design found. The exact synthesis design for (0×0016) is shown. Highlighted by the red and green colors is the partitioning of this circuit into two cells requiring only 5 and 7 gates and 1 DSM. The alternative design (not shown) given by the disjunctive form approach requires a higher requirement of 3 cells and 2 DSMs.

#### A 6-output 4-input design example

We consider a circuit with two 2-bit binary numbers *y*_1_*y*_0_ and *x*_1_*x*_0_ as inputs. As an example, let us design a full adder (*c*_2_*c*_1_*c*_0_), a subtractor (*d*_1_*d*_0_) and a comparator. The outputs are defined as: (*c*_2_*c*_1_*c*_0_) = (*y*_1_*y*_0_) + (*x*_1_*x*_0_), (*d*_1_*d*_0_) = |(*x*_1_*x*_0_) − (*y*_1_*y*_0_)|, where addition and subtraction here refer to the standard operations on the field of real numbers. The comparators are defined as follows: *e* = 1 iff (*y*_1_*y*_0_) = (*x*_1_*x*_0_) (and *e* = 0 otherwise), and *g* = 1 iff (*y*_1_*y*_0_) > (*x*_1_*x*_0_) (and *g* = 0 otherwise). A distributed design that shares minterms between the different BFs is shown in Figure 10.

**Figure 10:**
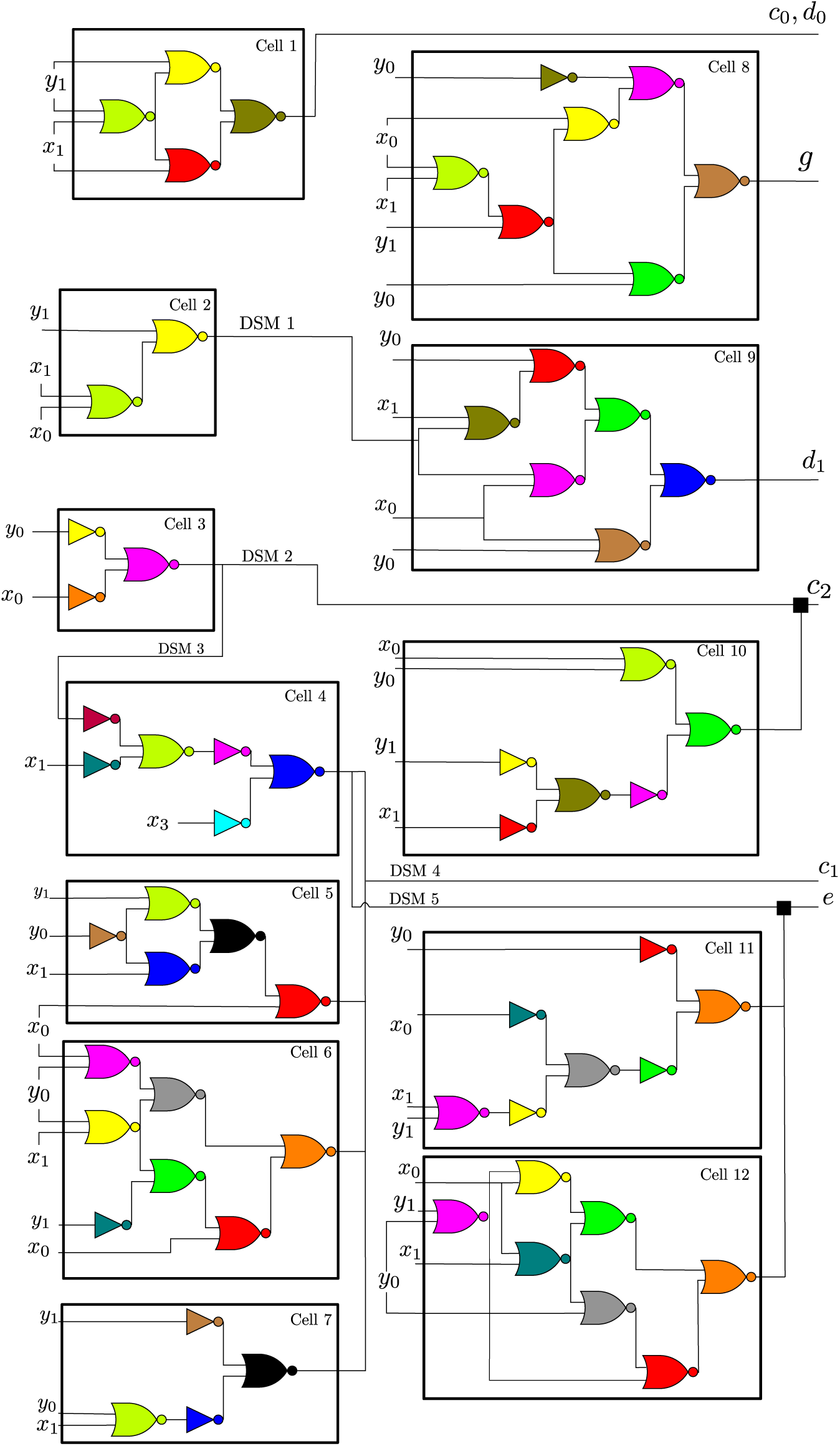
A 6-output 4-input circuit that uses 5 DSMs. The circuit implements a 2-bit adder, a 2-bit subtractor and a comparator. The outputs are defined as: (*c*_2_*c*_1_*c*_0_) = (*y*_1_*y*_0_) + (*x*_1_*x*_0_), (*d*_1_*d*_0_) = |(*x*_1_*x*_0_) − (*y*_1_*y*_0_)|, where addition and subtraction here refer to the standard operations on the field of real numbers. The comparators are defined as follows: *e* = 1 iff (*y*_1_*y*_0_) = (*x*_1_*x*_0_), and *g* = 1 iff (*y*_1_*y*_0_) > (*x*_1_*x*_0_). The symbol “■” refers to a DSM sensor.

## Discussion

Algorithms for implemented distributed genetic computation have been proposed. Taking in an BF *f* mapping and a given limit on the maximum number of NOR gates per cell *N*, a realization is calculated that fulfills the constraints. In particular, designs were provided for all 4-input BFs when *N* is 7 and the maximum number of DSMs *q* was 3. Combining different minterms to find an optimized disjunctive-form design leads a large reduction in the number of required cells. A database with results, useful for look-up when implementing circuits, has been made publicly available.

## Methods

### Background on Boolean Algebra

A Boolean function (BF) *f* of *n* inputs is a mapping *f*: {0, 1}^*n*^ → {0, 1}. Hence, there are 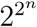 BFs with *n* inputs. A BF can be represented by at least three equivalent formalisms:

1. By specifying its truth table, i.e., listing its values (zero or one) for each of the 2^*n*^ possible binary vectors of *n* inputs. This list can be written as a string of 2^*n*^ binary digits. Thus, when there are *n* = 4 inputs, we specify functions as a string of length 16. For example, the four-input function 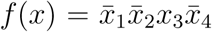 is nonzero when the input vector is (0, 0, 1, 0), and is zero otherwise. Since (0, 0, 1, 0) is the third vector in the list of all 16 possible vectors of length 4, the function would be specified as 00 … 00100, with a ‘1’ only in the 3rd entry from the end. A convenient shorthand is to use a string of ⌈2^*n*−2^⌉ hexadecimal digits (so, four hex digits when *n* = 4). The example above would be described as 0×0003 (the “0x” prefix indicates the use of hex notations).
2. By writing a Boolean algebraic expression that gives the value for each choice of inputs.
3. By a providing circuit *realization* that shows the interconnections between the gates.

There are two dual standard Boolean algebraic representations of BFs: the disjunctive and the conjunctive forms. Here, the disjunctive form (DF) is considered [30] which uses three Boolean operators: OR, AND, and NOT. For two Boolean variables *x, y* we use the following notation: *x* + *y*:= OR(*x, y*)*, xy*:= AND(*x, y*), and 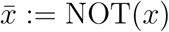. We also denote 1:= TRUE, 0:= FALSE, *x*^1^:= *x, x*^0^:= 1, and 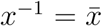. With these notations, any DF can be written as follows:

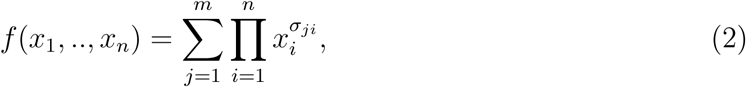

where *m* is the number of terms in the disjunctive form and *σ*_*ji*_ ∈ {1, 0, −1}.

A *literal* is a Boolean variable that is uncomplemented or complemented, e.g *x*_*i*_, 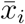 are literals. We say that a BF *c* is a *product term* if it is a product of literals. A product term is called a *minterm* of *n* variables if it has exactly *n* literals. Hence, a product term 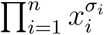 is a minterm if *σ*_*i*_ ∈ {±1}, *i* = 1,.., *n*. A minterm *c* can also be defined as a BF for which there exists a unique choice of Boolean variables 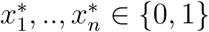 such that *c*(*x*_1_,.., *x*_*n*_) = 1 iff 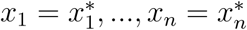. Hence, it follows that there are 2^*n*^ minterms. A DF of a BF *f* that is a sum of minterms is called a *disjunctive normal form* (DNF). It follows that the number of terms in a DNF is the number of *ones* in the truth table. The set of minterms that appear in the DNF of *f* is denoted by DNF(*f*).

It is often the case that the DNF can be reduced to a smaller number of terms. For example, the DNF 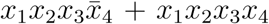 can also be written as simply *x*_1_*x*_2_*x*_3_. This is called the problem of DNF minimization. A reduction can be achieved by finding the so-called *prime implicants* of *f*. A product term *c* is called an implicant of *f* if the following statement holds: *c*(*x*_1_,.., *x*_*n*_) = 1 implies *f* (*x*_1_,.., *x*_*n*_) = 1. If the removal of any literal from an implicant *c* makes it a non-implicant, then *c* is called a *prime implicant*. We denote the set of prime implicant product terms of *f* by PI(*f*). The sum of all prime implicants is called the Blake canonical form. It can be reduced further by determining the *essential* prime implicants. A *minimized disjunctive form* (MDF) is a DF in which the number of terms is minimal. Computing an MDF can be handled by the Quine–McCluskey algorithm which can be used for a small number of inputs. For a larger number of inputs, the Espresso heuristic minimizer is the most popular algorithm [35]. Details of these algorithms can be found in [37].

A Boolean function which maps every element from {0, 1}^*n*^ to a value of 0 or 1 is called completely specified. For purposes of the algorithm to be discussed, it is also useful to introduce the notion of an “incompletely specified” or *partial* Boolean function, which maps a (possibly proper) subset of the elements from {0, 1}^*n*^ to a value of 0 or 1. An incompletely specified Boolean function *f* on *n* variables is a mapping *f*: *D* → {0, 1} where *D* ⊂ {0, 1}^*n*^. The set {0, 1}^*n*^\*D* gives the don’t-care conditions of the function *f*.

A (partial) Boolean function can be described using three sets: the on-set, off-set and don’t-care set. These three sets form a partition of the domain {0, 1}^*n*^ and can be used to fully describe the function in terms of the entire input space. The on-set of a Boolean function *f*, denoted on(*f*), is the subset of {0, 1}^*n*^ which *f* maps to 1. The off-set of a Boolean function *f*, denoted off(*f*), is the subset of {0, 1}^*n*^ which *f* maps to 0. The don’t-care set of a Boolean function *f*, denoted *DC*(*f*), is the subset of {0, 1}^*n*^ which *f* neither maps to 1 nor 0. It is the set {0, 1}^*n*^\*D*. Since the on-, off-, and don’t-care sets form a partition of the input space, only two of the three sets are required in order to describe the function. Using the on- and off-set notation we can distinguish between completely-specified and incompletely-specified functions. A completely-specified function has an empty don’t-care set while the don’t-care set of an incompletely specified function is non-empty.

If *L* and *U* are Boolean functions, then the interval [*L, U*] is the set of functions defined by [*L, U*] = {*f ∈ B*: *L ≤ f ≤ U*}, where *B* is set of all *n*-variable completely specified Boolean functions. *L* is the lower bound of the interval, while *U* is the upper bound of the interval. The requirement *L ≤ f* implies that every function *f* contained in the interval must evaluate to 1 on the same set of input combinations on which *L* evaluates to 1. The requirement *f ≤ U* implies that every function *f* in the interval must evaluate to 0 for the same set of input combinations on which *U* evaluates to 0. Intervals can be used to describe incompletely specified functions. The interval [*L, U*] represents the Boolean function that evaluates to 1 on the same set of input combinations as *L* and evaluates to 0 on the same set of input combinations as *U*. The incompletely specified function function *f* can be defined [⌈*f1⌉*, ⌈*f*⌉]. The relationships between the upper and lower bounds of a Boolean function *f* with the on-, off-, and don’t-care sets of *f* are as follows:

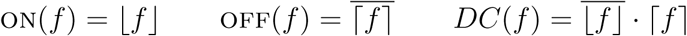

### Formal definition of the problem

An (*N, q*)-network is a realization of an *n*-input BF mappings *f*_1_,.., *f*_*p*_, represented as a directed acyclic bipartite graph (also known as a acyclic Petri net [38]), for which:

1. “places” correspond to cells (the output node is a cell with no out-edges);
2. “transitions” correspond to *QS nodes* or *input nodes*. The input node has no in-edges.

Assume there are *m* cells, each cell *i* computes a BF *c*_*i*_, *i* = 1,.., *m* that is realizable by a maximum of *N* NOR2 gates. A QS node then computes an OR gate of these outputs.

#### Definition 1.

*Let f*_*j*_: {0, 1}^*n*^ → {0, 1}, *j* = 1,.., *p be BFs. The vector BF F*:= (*f*_1_,.., *f*_*p*_) *is said to be* (*N, q*)*-realizable if there exists an* (*N, q*)*-network, with m cells and q QS nodes, and BFs c*_1_,..., *c*_*m*_, *each of which is realizable by at most N NOR2 gates. The function F is evaluated by the network as follows: each cell computes the corresponding function c_i_ and each QS node computes an OR operation. For each input w* ∈ {0, 1}^*n*^, *w*_*i*_ *is substituted into the ith input node of the network and it is used to compute values for all cells and QS nodes. The value of the jth output node is taken as the value of f*_*j*_(*w*).

**Remark 1.** *Instead of distinguishing between the two types of node: cells and QS, an equivalent graphical description is in terms of directed hypergraphs. A directed hypergraph is specified by a set of nodes, which can be taken as the cells, and a set of “hyperedges”, i.e. pairs* (*S_i_, T_i_*) *where each of S_i_ and T_i_ is a subset of the set of cells, thought of as the “source” and “target” of the “hyperedge” in question. Each QS node may be replaced by a hyperedge that has as its source the in-edges to QS and as its target the out-edges of QS. However, it is intuitively preferrable to use explicitly the two types of nodes as each of them computes a Boolean function in our application. Physically, one may think of QS nodes as concentrations of quorum-sensing signals in the environment of the cell population, making the mapping into physical variables more transparent*.

### Exact synthesis algorithm for (*N*, 0)-networks

The next step is to find the circuit representation using a minimum number of NOR2 gates for each cell in the design. The exact synthesis algorithm described here does this by systematically generating circuits in search of the one that uses the minimum number of NOR2 gates. The algorithm represents a group of circuits using a partial network. The circuits represented by a partial network are those that can be created by adding edges and possibly nodes to the partial network. The algorithm searches through the space of possible circuits by incrementally completing the partial network. The search space is divided by choices made when completing the partial network. This results in a branch-and-bound algorithm that recursively performs an exhaustive search of the space in order to guarantee the optimality of the solution. To make the exhaustive search more efficient, the algorithm makes use of pruning techniques to help minimize the search space.

#### Networks and Boolean Functions

The Boolean function associated with a node in a network is the function produced at the output of this node. The function can be expressed either in terms of its immediate inputs or in terms of the primary inputs of the circuit. The local function is the Boolean function associated with the gate expressed in terms of its immediate inputs (NOR2), while the global function is the function associated with the gate expressed in terms of the primary inputs.

A relationship exists between the global functions at the inputs and output of a gate node based on the local function of the node. The relationship for the NOR2 gate is as follows:

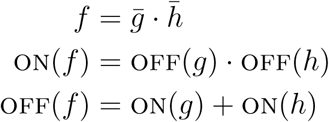

During intermediate stages of the algorithm partial or incomplete networks will be created. A partial network can be viewed as representing a set of circuits, namely those that can be created by adding edges and possibly nodes to the partial network. There are two types of gate nodes in a partial network, bounded and unbounded nodes. A node of a partial network is bounded if the size of the fan-in set has reached 2, the fan-in restriction imposed by using NOR2. A node is unbounded if the size of the fan-in set is less than 2. The Boolean function of a gate node will be determined by inputs to the gate. However, since the network is incomplete, there is a possibility that some nodes may be unbounded. The unassigned inputs of an unbounded gate node can take any function. Therefore when computing the global function of an unbounded node, the unassigned inputs are given the function [0, 1] (ON = 0, OFF = 0).

The algorithm moves through the search space by adding nodes and edges to the partial network. The addition of an edge or a node to a partial network will have an impact on the function associated with the nodes on either side of this edge or node. In addition, the change in the function of a node may have an impact on the function of the nodes in its fan-in and fan-out sets. The relationships between the inputs and outputs of the NOR2 gate given above will be used to determine the implications that result when changes are made in a partial network. These implications will keep the global functions at the inputs and outputs of each gate node consistent with the local function described by the logic gate.

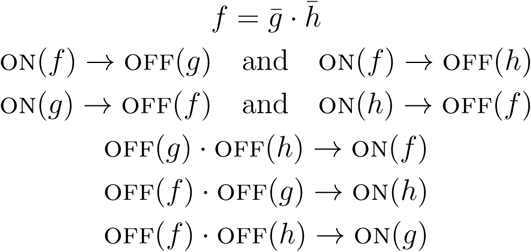

#### Connectible Nodes

There are two sets of constraints which determine what edges can be added to a partial network. These constraints are based on the functional and structural properties of the network. The functional constraints determine which nodes can be connected to the input of a given gate node based on the global functions of the nodes. Given the current function at a node N and the functions at its inputs, a constraint function can be created which when solved will give the permissible functions for the node’s inputs. Any node whose Boolean function falls within this set of permissible functions will be considered functionally connectible to the node N. The first step is to find the constraints for the set of permissible functions. For a NOR2 node, a function g is permissible as an input to a node if 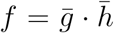 where *f* is the function at the output of the node and *h* is the function at one of the inputs to the node. If *f* and *h* are completely specified functions, then any function *g* that satisfies this constraint would be considered a permissible function as an input to the node. However, the functions *f* and *h* are not always completely specified.

The constraint for determining the permissible functions must allow for incompletely specified functions at both *f* and *h*. A function *g* is permissible as an input to a node with output function [*⌈f⌉, ⌈f⌉] and input function [*⌈h⌉*, ⌈h⌉*] if for some function *f* ∈ [*⌈f⌉*, ⌈*f*⌉] and some function *h* ∈ [*⌈h⌉*, ⌈*h⌉*], 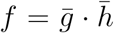. This can be simplified into the following constraint.

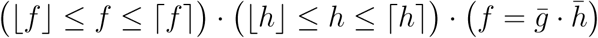

Any function *g* which satisfies this constraint is considered a permissible function to the node. There may be more than one function that satisfies this constraint. The interval describing the set of permissible functions can be found directly after some transformations on this constraint.

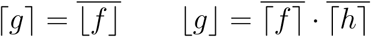

A node *C* in a network is functionally connectible to a gate node *N* if the intersection of *C*’s function interval with the interval of permissible functions for an input of *N* is nonempty. The intersection of the two intervals gives the set of functions from the permissible set that the node *C* can take. Let *C* ∈ [⌈*C*⌉, ⌈*C*⌉] be the function at node *C*. *C* is functionally connectible to a gate node *N* with output function *f*, input function *h*, and permissible functions *g* if the following condition is satisfied:

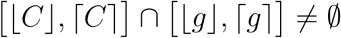

This requirement can also be stated as:

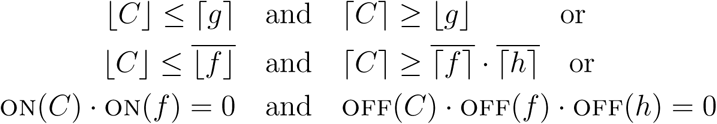

The second set of constraints that must be satisfied when adding edges to the network are structural constraints. These constraints are needed to insure that the network remains acyclic. A node C is structurally connectible to a node N if C does not appear on any path from N to a primary output. In other words, C can not be an ancestor of N.

The connectible set of a node N is the set of nodes in the partial network that are both functionally and structurally connectible to N.

#### Covering

The notion of covering is central to the exact synthesis algorithm. A minterm in the off-set of the global function of a NOR2 gate node *N* is covered if the minterm appears in the on-set of the global function of at least one of *N*’s fan-in nodes. Using this notion of covering, the off-set minterms of a NOR2 gate can be divided into two sets. The covered set is the set of minterms that are already covered by a fan-in of the node: OFF(*f*) ON(*g*)+ON(*h*), where *f* is the global function describing the output of the node and *g* and *h* are the global functions describing the two inputs to the node. The uncovered set is the set of minterms that are not yet covered by the inputs of the gate node: 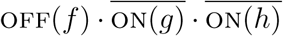. When the uncovered set becomes empty, the node is completely implemented by its inputs. These nodes are called covered. The algorithm will perform a covering when the fan-in set of a node is changed or the global function of one of the fan-in nodes is changed so that at least one minterm from the node’s uncovered set moves to its covered set.

#### Algorithm Description

The branch-and-bound algorithm will explore the search space recursively. Each step of the recursion will perform a covering of at least one uncovered minterm from some node in the partial network. When the current network is determined to be complete or its cost becomes larger than the current cost bound, the recursion stops and backtracks to the previous step.

The algorithm is initially called on a partial network that represents the entire set of circuits in the search space. This initial network will contain one gate node for each output function and one input node for each primary input. The fan-in and fan-out sets of all these nodes are empty, while the global function at each node is set according to the function it represents. In addition, the connectible set for each of the gate nodes must be computed according to the constraints described above.

The ExploreNetwork procedure is the main procedure of the algorithm. It takes as input a single partial network. It performs a covering of some uncovered minterm in the partial network and then recursively call the same procedure using the new partial network. The pseudocode for the ExploreNetwork procedure is given in below.

##### Algorithm 1: Exact Synthesis Algorithm

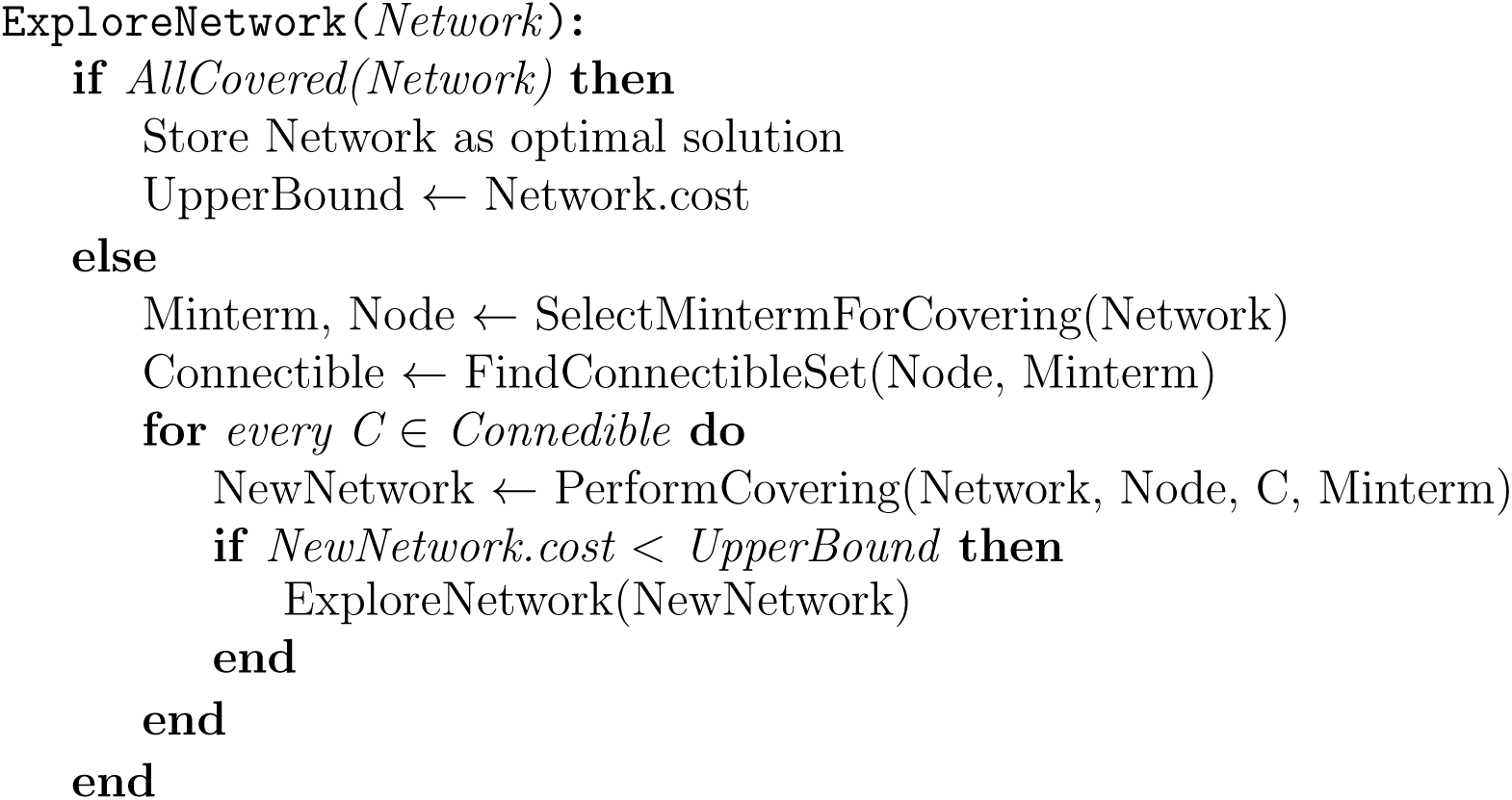

The *ExploreNetwork* procedure begins by first determining whether every gate node in the network is covered by its inputs. This is performed by the *AllCovered* function. A gate node *G* is covered by its inputs *I*1 and *I*2 if OFF(*G*) = ON(*I*1) + ON(*I*2). If every gate node is completely covered,the network is a complete circuit. Therefore it can be stored as the current optimal network and the cost bound can be set accordingly.

If there are nodes remaining in the network that are not fully covered by their inputs, then work still remains towards completing this partial network. The procedure will attempt to cover an uncovered minterm from at least one node. The covering process begins by first selecting an uncovered minterm and its corresponding node from the network. This is done by the *SelectMintermForCovering* procedure. The minterm that is chosen by this procedure is guaranteed to be covered during this step of the recursion. It may be the case that additional uncovered minterms in the network may be covered as well. This will depend on the covering that is made.

The procedure continues from the minterm selection to obtaining the connectible set for the node whose off-set minterm is to be covered. A connectible set is stored for every node. This set gives all the possible ways that exist in the network for covering every uncovered minterms from the node. The procedure *FindConnectibleSet* will prune the node’s connectible set down to only those nodes that can cover the selected minterm. In order for a node *C* in *Node*’s connectible set to be included in *Connectible*, the minterm *Minterm* must not appear in the off-set of *C*. *C* is added to *Connectible* if off(*C*) *Minterm* = 0. A node *C* satisfying this condition can be used to cover *Minterm* since *Minterm* must be contained in (or moved to) *C*’s on-set in order for *C* to cover *Minterm*. The *FindConnectibleSet* procedure will also add a new gate node to the set *Connectible* if the fan-in set of *Node* is less than 2. This gate node will be a new node which does not yet appear in the network.

Each node in the connectible set returned by the *FindConnectibleSet* procedure must be considered as a way of covering the selected minterm. The enumeration of the possible coverings using each node in the connectible set corresponds to the branching part of the branch-and-bound search. For a single connectible node, *C*, a new network is created by the *PerformCovering* procedure. This new network will be a supernetwork of *Network* with the added covering of *Minterm* using *C*.

The process of covering *Minterm* can add a connection between the nodes *Node* and *C* as well as change the global function at *C*. Based on the relationship between the global functions at a node’s fan-in and fan-out described by the node’s local function, the changes made at *Node* and *C* can effect the global functions at other nodes in the network. The changes to the global functions that result from a covering are called functional implications of the covering. These functional implications must be completed to maintain the consistency of the global functions.

Once the functional implications have been completed, the *PerformCovering* procedure updates the connectible set of each node in the network. Since the global function may have changed for some nodes, the set of nodes that are now connectible may have also changed. It is possible that previously covered nodes have now become uncovered, and therefore the set of connectible nodes will need to be computed for these nodes as well. The connectible set of each uncovered gate node in the network can be found according to the constraints described above. The connectible set update will conclude the *PerformCovering* procedure.

Once the covering is complete, the cost of the partial network is compared to the current cost bound. If the cost of the new partial network is greater than the current cost bound, this network can be pruned. This is the case since each elementary network in the set that the new network represents will have cost larger than the current cost bound. Therefore, none of the elementary networks represented can be optimal. If the cost of the new network is less than or equal to the cost bound then it is possible that some elementary networks in the set that this partial network represents may have cost less than or equal to the current cost bound. Thus requiring this portion of the search space to be explored further. The portion of the search space represented by this new partial network is explored by calling the *ExploreNetwork* procedure on the new network.

When the recursive call returns, the next node in the connectible set is used to cover the selected minterm. Once again the search space represented by the new partial network is explored. Each possible way of covering the minterm is selected and then explored until all options have been exhausted. Once the recursion returns for the last time, all complete networks represented by the original partial network have been explored. Any optimal networks represented by the original partial network have been stored as such. Now the procedure can return to the previous recursive call since this call has been completed.

### Minterms: realizable and offending

Our framework proposed in Fig. 2 relies on representing a BF as a disjunction of other BFs. Hence, we first study the DNF in terms of minterms. We provide the following definition:

#### Definition 2.

*Let N be given. A minterm in n variables is said to be <u>offending</u> with respect to N if it cannot be realized with N NOR2 gates or less. The set of all offending minterms with respect to N is denoted as 𝒪_N_*.

The following statement that can be proven using various methods including the exact synthesis algorithm which we have introduced earlier.

#### Theorem 2.

*Let c be a minterm of n variables with n ≥ 2. Then the minimum number of NOR2 gates to realize c is* 3(*n* − 1) − *q*(*c*), *where q*(*c*) *is the number of complemented literals in c*

**Example.** The minterm *x*_1_*x*_2_ needs (3)(1) – 0 = 3 gates, while the minterm 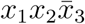 needs (3)(2)-1=5 gates.

It follows that any minterm of *three* variables or less can be realized with six gates or less. Therefore, the design in Figure 2 can realized *any DF* of any BF *f* of three variables or less.

For minterms with four variables, it follows that a minterm with four or three uncomplemented literals cannot be realized with 7 NOR gates. The minterm *x*_1_*x*_2_*x*_3_*x*_4_ needs at least 9 NOR gates, and the minterms 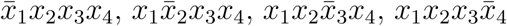 need at least eight NOR gates. These five minterms are called the *offending minterms* with respect to *N* = 7 gates. If a disjunctive-form design contains one or more offending minterms, additional DSMs are used to divide up these circuits further.

### Lower bound on the number of (*N*, 1)-realizable BFs

We state the following Theorem:

#### Theorem 3.

*Let f be a BF, and let N be given. If 𝒪_N_ ∩PI(*f*) = ∅, then f is (*N*, 1)-realizable*.

*Proof.* Let us write *f* as an 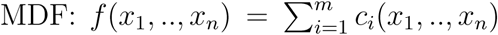. By the assumption, *c*_*i*_ ∉ 𝒪_N_ for all *i*. Hence, *f* can be realized using the disjunctive form design in Figure 2 where each *c*_*i*_ is (*N*, 0) realizable.□

In other words, any function that does not have a prime implicant offending minterm is (*N*, 1)-realizable. Hence, our objective in this subsection is to count those BFs combinatorially.

In order to facilitate the discussion, a binary representation of minterms is used. An *n*-variable minterm can be represented with *n* bits. If a literal is uncomplemented then it is represented as 1, otherwise it is represented as 0. For instance, the minterm 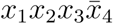 is written as 1110. Using this notation, the *distance* between two minterms is defined as the *Hamming distance* between their binary representations. Two minterms are *neighbors* if their distance is 1. Hence, we are ready to state the following result:

#### Lemma 4.

*Let f be a BF, and let c ∈ DNF*(*f*) *be a minterm. If there exists a minterm c^∗^ ∈ DNF*(*f*) *such that c, c^∗^ are neighbours then c, c^∗^ are not prime implicants*.

*Proof.* Since both *c, c^∗^* DNF(*f*), then *c* + *c^∗^* appears in the DNF of *f*. But since *c, c^∗^* differ in only one literal, then *c*+*c^∗^* simplifies into a product term with a smaller number of literals. The resulting product term is an implicant of *f*, hence *c, c^∗^* are not prime implicants. □

The following test follows:

#### Theorem 5.

*A BF f is* (*N*, 1)*-realizable if for every c ∈ 𝒪*_*N*_ ∩ *DNF*(*f*), *there exists c*^∗^ *DNF*(*f*) *such that c, c^∗^ are neighbors*.

Hence, our task reduces to counting BFs that have a minterm in 𝒪_*N*_ ∩ DNF(*f*) without neighbours. We propose a fast algorithm to find the set of BFs that satisfy the conditions of Theorem 5. We use the notation 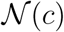 as the set of neighbors of a minterm *c*. The algorithm is given below:

**Parameters:** *n* number of variables, *N* the maximum number of gates per cell.

**Initialization:** Set *X* = 0;

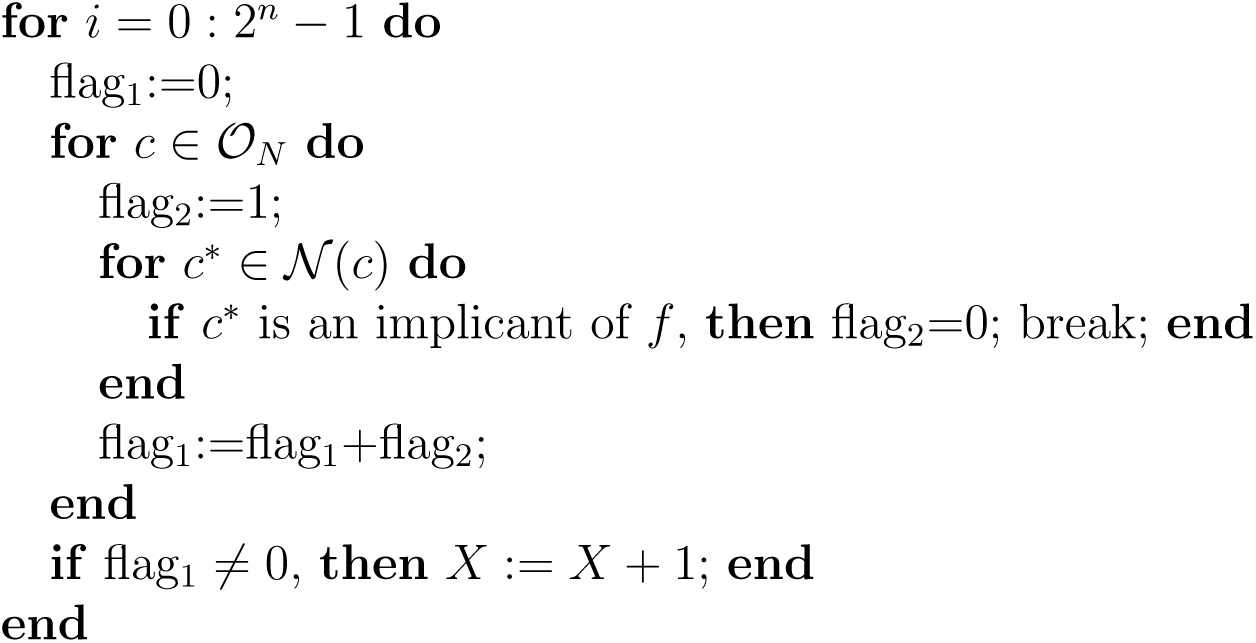

**Output:** *X* is an upper-bound on the number of networks which are *not* (*N*, 1)-realizable.

#### Realization with offending minterms: Proof of Theorem 1

We provide a constructive proof of Theorem 1. If 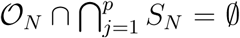, then the numbers of DSMs needed is *p* at most. Next, we study the cases of more than one offending minterm. Remember that *𝒪*_7_ = {1111, 1110, 1101, 1011, 0111} and *𝒪*_8_ = {1111}. For the *N* = 9, we have *𝒪*_9_ = *∅*, and hence the corresponding statement follows directly.

##### Vector BFs with one offending minterm

We study the case of only one offending minterm appearing as a prime implicant. For instance, consider the minterm 1111 which is offending for both the cases *N* = 7 and *N* = 8.

Let 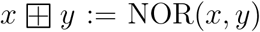, and remember that 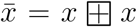. A minimal realization of the minterm 1111 consists of 9 gates and is given as follows:

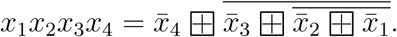

One DSM molecule can be used to partition the realization above into two cells with four and five gates as follows:

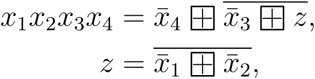

where *z* is communicated via a DSM. In function format, we are writing *x*_1_*x*_2_*x*_3_*x*_4_ = *f*_1_(*f*_2_(*x*_1_*, x*_2_)*, x*_3_*, x*_4_) where *f*_2_ is (*N*, 0)-realizable and *f*_1_ is (*N*, 1) realizable, where *N* ≥ 5. This partition is depicted in Figure 3-a). Note that this completes the proof of the statement of Theorem 1 for the case *N* = 8.

For the case *N* = 7. The other four offending minterms can be treated similarly. Their realizations requiring 8 NOR gates can be partitioned into two 4-gate cells.

##### BFs with two offending minterms

For any two minterms chosen, they will share exactly two uncomplemented literals. For instance consider 1110, 1101. Both *x*_1_*, x*_2_ are uncomplemented. Then, the following realization is proposed:

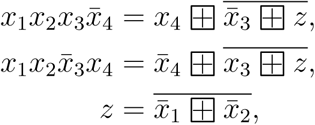

where *z* is communicated via a DSM. Hence, we compute *x*_1_*x*_2_, and then use it for computing the full minterm. This partition is depicted in Figure 3-a).

Hence, any vector BF with no more than two offending minterms needs no more than *p* + 1 DSMs.

##### BFs with three or more offending minterms

If three or more offending minterms in the set {1110, 1101, 1011, 0111, 1111} appear in the MDFs of a vector BF then we will show below that no more 2 DSMs are needed.

Let’s consider the case of five offending minterms. It can be seen that this case can be realized with two DSMs as follows. The four minterms can be divided in no particular order into two subsets which each share exactly two uncomplemented literals. Then, a similar design follows.

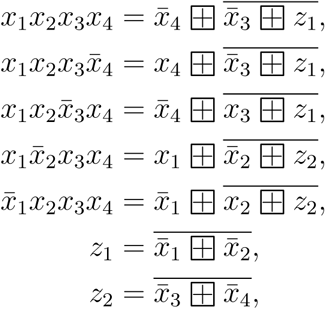

where *z*_1_ and *z*_2_ are communicated via two different DSMs. Hence, no more than *p* + 2 DSMs are needed.

### Counting (7, 2) and (7, 3) realizable BFs

In the case of a scalar BF, we can count the number of BFs realizable with the disjunctive form design combinatorially. For a single offending minterm. We note that if 1111 ∈ DNF(*f*), then the rest of the offending minterms cannot appear, otherwise, they will be neighbours, and the associated BF will be (7,1)-realizable. This amounts to specifying a fixed value to the vector (*f* (1, 1, 1, 1), *f* (1, 1, 1, 0), *f* (1, 1, 0, 1), *f* (1, 0, 1, 1), *f* (0, 1, 1, 1)), namely (1, 0, 0, 0, 0) out of 2^5^ possible choices. The total number of these BFs is 2^16^/2^5^ = 2^11^ = 2048. The other four minterms need be handled together since they share neighbors. Similar combinatorial computations gives the total number of BFs with one offending minterm to 7,808.

If two offending minterms in the set {1110, 1101, 1011, 0111} appear in the DNF without neighbours then these have been shown to be (7, 2)-realizable. The total number of BFs with exactly two offending minterms is 960.

The total number of BFs with exactly three offending minterms is 2^7^ = 128 BFs. For the case of a scalar BF we can do better than the upper bound given in Theorem 1 since any three offending minterms can be realized with one DSM as follows:

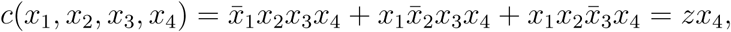

where 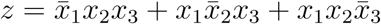 can be computed via a single DSM.

Hence this expands the set (7, 2) scalar BFs to 8896 using the disjunctive design. Finally, there are 2^5^=32 BFs with four offending minterms.

## A graph partitioning algorithm

This algorithm is implemented by first expanding the design provided by the exact synthesis algorithm so that the output of each NOR2 gate is only used once, effectively creating repeated ‘branches’. For each gate in the expanded design, the total number of gates is computed such that this number accounts for all the gates located downstream of a given gate plus the gate itself. To determine that a design can be partitioned using 1 DSM, we first look at the total number of gates of the upstream node *T*_*up*_. Going through the other gates, there must exist one gate *i* such that its total number of gates *T*_*i*_ with *n* occurences within the expanded design that fulfills the following conditions:

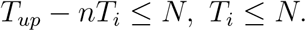

In the case of partitioning using 2 DSMs, two gates *i* and *j* are used. There are two possibilities of partitioning, one in which neither *i* or *j* is located downstream (or upstream) of the other gate. In this case, the conditions are simply that:

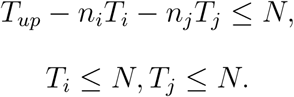

The second scenario occurs when gate *i* is either located upstream or downstream of *j*. By assuming that we order *i* and *j* such that *i* is the upstream gate, the conditions to be realizable with 2 DSMs become:

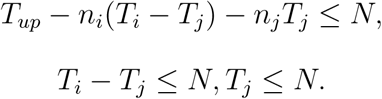

## Acknowledgments

This work was supported by NSF grant 1849588 and SRC grant SB-2837-B.

